# Accessory B Chromosomes Alter Chromatin Accessibility and Gene Expression in Maize Pollen Nuclei

**DOI:** 10.64898/2026.11.09.765325

**Authors:** Petr Urbiš, Lucie Hloušková, Kateřina Holušová, Jaroslav Doležel, Jan Bartoš

**Affiliations:** Institute of Experimental Botany of the Czech Academy of Sciences, Centre of Plant Structural and Functional Genomics, Šlechtitelů 31, Olomouc, 779 00, Czech Republic; Department of Cell Biology and Genetics, Palacký University, Olomouc, Czech Republic

## Abstract

B chromosomes in maize (*Zea mays* L.) are supernumerary, nonessential chromosomes that exhibit non-Mendelian inheritance through chromosome drive. During the second pollen mitosis, they undergo nondisjunction, resulting in their preferential transmission via B-containing sperm cells. Although their transmission mechanisms have been extensively characterized, the effects on chromatin organization and transcriptional regulation in gametophytic cells remain poorly understood. Here, we combined flow cytometric sorting of pollen nuclei with an assay for transposase-accessible chromatin sequencing (ATAC-seq) and RNA sequencing (RNA-seq) to investigate the impact of B chromosomes on chromatin accessibility and gene expression in mature maize pollen. Using pollen nuclei isolated from a maize inbred line B73 carrying zero and two B chromosomes, we generated high-purity datasets suitable for integrated chromatin and transcriptomic analyses. We identified extensive chromatin accessibility and transcriptional activity at B-chromosome loci. Importantly, the presence of B chromosomes was also associated with altered chromatin accessibility and gene expression on the standard A chromosomes, revealing regulatory effects extending beyond the B chromosome itself. Together, these findings contribute to a broader understanding of effects of selfish genetic elements in plants.

**Teaser:** B chromosomes contain functional regulatory elements that drive active gene expression in mature maize pollen.

## Introduction

Pollen grains, the microscopic carriers of genetic material in plants, play a crucial role in the reproductive process of angiosperms and gymnosperms and are vital for the continuation of plant species. Consequently, isolating and studying the male gametophyte—a key factor in the evolutionary success of flowering plants and a uniquely accessible cell type—is thus of great interest (Honys and Twell, 2003; Chumak, 2015). To facilitate their study, flow cytometry (FCM) has been found useful, despite the challenges posed by the chemically inert and physically resilient pollen cell wall, which severely hinders the mechanical disruption required to release intact nuclei (Chumak, 2015; Kron *et al*., 2021). Competition between male gametophytes for the fertilization of a limited number of egg cells is common in sexual reproduction, and pronounced differences in pollen tube growth rates reflect genetic differences among individual microgametophytes (Mulcahy, 1979; Mulcahy *et al*., 1996; Honys and Twell, 2003; Chumak, 2015). At the molecular level, these functional differences are increasingly linked to epigenetic landscapes.

In addition to the standard karyotype, widespread populations of plant, animal and fungal species contain nonessential, dispensable, supernumerary (i.e., “extra”) B chromosomes (Longley 1927). These chromosomes are often preferentially inherited and deviate from the Mendelian law of equal segregation via a process known as chromosome drive (Carlson *et al*., 2006; Wu *et al*., 2019). In maize, the persistence of the B chromosomes is governed by a two-component drive mechanism: nondisjunction at the second pollen mitosis (Roman, 1947; Brennan *et al*., 2023) and preferential fertilization of the egg cell by the B-containing sperm during double fertilization (Roman, 1948; Brennan *et al*., 2023). During the second pollen mitosis, the sister chromatids of the B chromosome fail to separate at the centromeric region. This nondisjunction occurs exclusively in the generative cell, yielding two sperm nuclei—one carrying two B chromosomes and the other carrying none—while the vegetative nucleus remains unaffected.

These B chromosomes have distinctive repeat-rich regions around the centromere (ZmBs repeats, knob-like heterochromatin, etc.), so the nondisjunction is controlled by cis and trans factors on the B chromosome (Birchler and Yang, 2021; Blavet *et al*., 2021). In pollen, B-specific DNA is often localized to a particular tip or subregion of the sperm nucleus, giving that sperm a unique spatial chromatin architecture that could influence how it interacts with the egg or polar nuclei during double fertilization (Rusche *et al*., 1997; Blavet *et al*., 2021). In preferential fertilization, the sperm harbouring two B chromosomes enters and fertilizes the egg cell at a higher frequency compared to the sperm without Bs (Roman, 1948; Carlson, 1969; Carlson and Phillips, 1986; Brennan *et al*., 2023), thus enhancing the transmission rate of B chromosomes into the embryo and subsequent progeny (Roman, 1948; Brennan *et al*., 2023).

Mechanistically, preferential fertilization by B-carrying sperm in maize seems to involve specific sequences or structural features near the centromere-proximal region of the B chromosome, alongside egg-cell-driven recognition of one sperm nucleus over the other, although the exact molecular players remain largely unknown (González-Sánchez *et al*., 2003; Brennan *et al*., 2023). Chromosome B-A translocation studies indicate that the determinant of preferential fertilization maps to a region at or very near the B centromere, implying the involvement of centromere-associated DNA, chromatin, or proteins (Beckett, 1991; Blavet *et al*., 2021). Furthermore, the B-centric heterochromatin contains a site required for nondisjunction (Carlson and Chou, 1981; Carlson *et al*., 2006). This suggests that local repeats or chromatin features may jointly promote both nondisjunction and biased fertilization, possibly via altered centromere/kinetochore behaviour or nuclear positioning (Rusche *et al*., 1997; Blavet *et al*., 2021). New developments in analyzing sequence composition provide a valuable opportunity to examine the underlying structure of B chromosomes. Under ordinary conditions, B chromosomes do not bestow any advantages on the organisms that harbour them but rather represent a specific type of selfish DNA (Jones and Rees, 1969; Carlson *et al*., 2006). The highest number of B chromosomes that a host species can bear is determined by the chromosomes’ origin and a balance between the B chromosome drive and the negative impacts on fitness, particularly reduced fertility when Bs are present in higher numbers (Bougourd and Jones, 1997; Wu *et al*., 2019).

In this study, we investigated the effect of B chromosomes on pollen nuclear chromatin accessibility and the transcriptome. By comparing ATAC-seq and RNA-seq datasets from flow-sorted nuclei of mature maize B73 pollen grains with and without two B chromosomes (+B/0B), we provide the first insights into the molecular behaviour of male gametophytes carrying these accessory chromosomes.

## Results

### RNA-seq

We performed RNA-seq on mature pollen of maize B73 (+B/0B) accessions in three biological replicates. We generated on average ∼35 million high-quality paired-end 150-bp reads (Supplementary Table S1) and mapped them to the *Zea may*s reference genome RefGen_v5 supplemented with the reference of the maize B chromosome (Hufford *et al*., 2021; Hloušková *et al*., 2025). Out of 40,445 annotated genes, 8,679 were expressed (TPM >1) in B-negative mature pollen (0B samples) and 9,462 in +B samples (Fig. 1A). Maize mature pollen grains exhibit an overall twofold lower gene-expression level and fewer expressed genes with TPM >1 than anthers and many other tissues (Shi *et al*., 2021; Han *et al*., 2022; Nelms and Walbot 2022; Zhang *et al*., 2022; Hloušková *et al*., 2025). However, they also contain a set of distinct pollen-specific genes, characterized by a profound enrichment in mature pollen compared to other types of tissue (Supplementary Dataset S1). We performed differential expression analysis to investigate +B pollen, with particular attention paid to the expression of B chromosome-encoded genes. Out of 689 predicted B chromosome-located genes, we identified 103 transcribed genes defined as having transcripts per million (TPM) value greater than 1 in mature pollen (Supplementary Dataset S2), 98 of these were shared with B chromosome-encoded genes expressed in mature anthers with TPM >1 (Hloušková *et al*., 2025).

**Figure 1.**
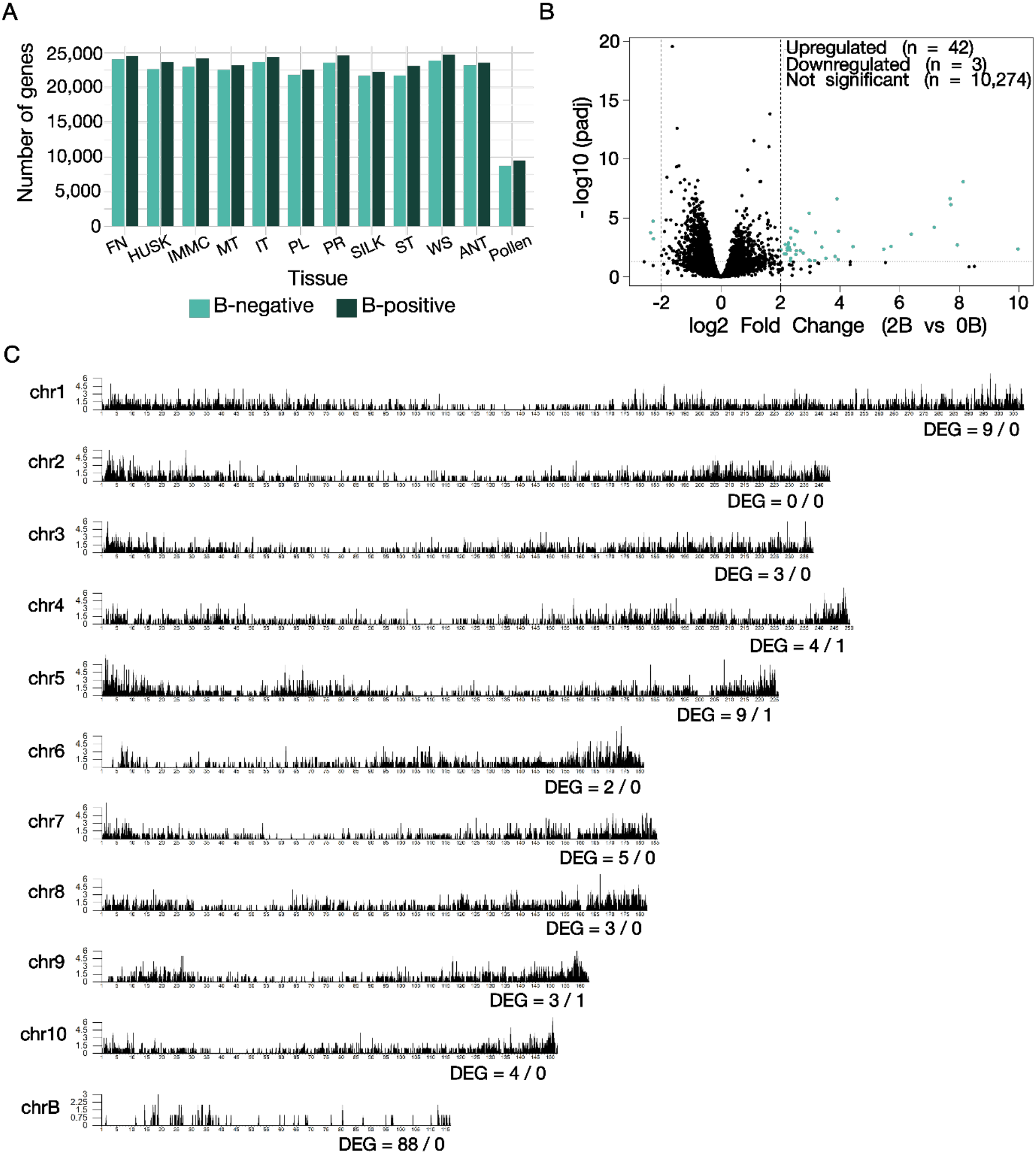
Gene expression and chromosomal distribution of expressed and differentially expressed genes in maize pollen. **(A)** Number of A-chromosome genes with TPM > 1 detected in different maize tissues. Tissues are: FN, first stem node; HUSK, husk; IMMC, unpollinated immature cob; MT, meiotic tassel; IT, immature tassel; PL, first leaves at the one-leaf stage (V1); PR, primary root; SILK, silk; ST, leaf tip of the youngest actively growing leaf at the five-leaf stage (V5); WS, whole germinated seedling; ANT, anthers with mature pollen; and Pollen, mature pollen. Data for all tissues with exception of Pollen were originally published by Hloušková *et al*. (2025). **(B)** Volcano plot shows differential gene expression between 2B and 0B pollen. The x-axis represents log2 fold change and the y-axis represents -log10 (adjusted *P* value). Dashed lines indicate the thresholds used to define differentially expressed genes. A total of 42 A-chromosome genes were upregulated and three genes were downregulated, while 10,274 genes were not significantly differentially expressed. **(C)** Distribution of expressed (TPM > 1) genes along chromosomes 1-10 and the B chromosome. Gene density is shown along each chromosome, and the numbers indicate the number of upregulated/downregulated differentially expressed genes identified on the respective chromosome.

### Differentially expressed genes

Analysis of differentially expressed genes (using DESeq2) identified 88 differentially expressed genes on the B chromosome and very low number (42) of upregulated and downregulated (3) genes encoded by A chromosomes in the presence of two B chromosomes (Figs. 1B; 1C). Expectedly, no downregulated B chromosome genes were detected by DESeq2 (Supplementary Dataset S3), as such category lacks biological meaning due to their complete absence in 0B samples. In comparison, in maize anther with mature pollen, the DESeq2 analysis previously detected 244 upregulated genes expressed on B chromosomes alongside 808 upregulated and 348 downregulated genes encoded by A chromosomes under the same condition (Hloušková *et al*., 2025). Our findings extend and refine the results of Hloušková *et al*. (2025; Supplementary Fig. S1), who reported that the expression of B-encoded genes is most prevalent in reproductive organs.

Subsequently, we performed a gene ontology (GO) enrichment analysis to investigate the functional roles of the 88 B chromosome- and 42 A chromosome-encoded upregulated genes. These genes were compared against all annotated A- and B-localized genes as the respective reference sets Where several significant terms were related, we reported the more specific term (i.e., the one lower in the GO hierarchy) over its more general parent term to maximize informativeness. The complete lists of tested GO terms, including non-significant ones, are provided in Supplementary Dataset S4.

Among biological processes, A-chromosome genes were significantly enriched for “gene expression” (GO:0010467), followed by “cellular response to stress” (GO:0033554) and “macromolecule biosynthetic process” (GO:0009059). In the molecular function category, the most enriched GO term was “ATP binding” (GO:0005524), followed by “ATP-dependent activity” (GO:0140657). In contrast, no significantly enriched GO terms were identified within the cellular component category (Fig. 2B).

**Figure 2.**
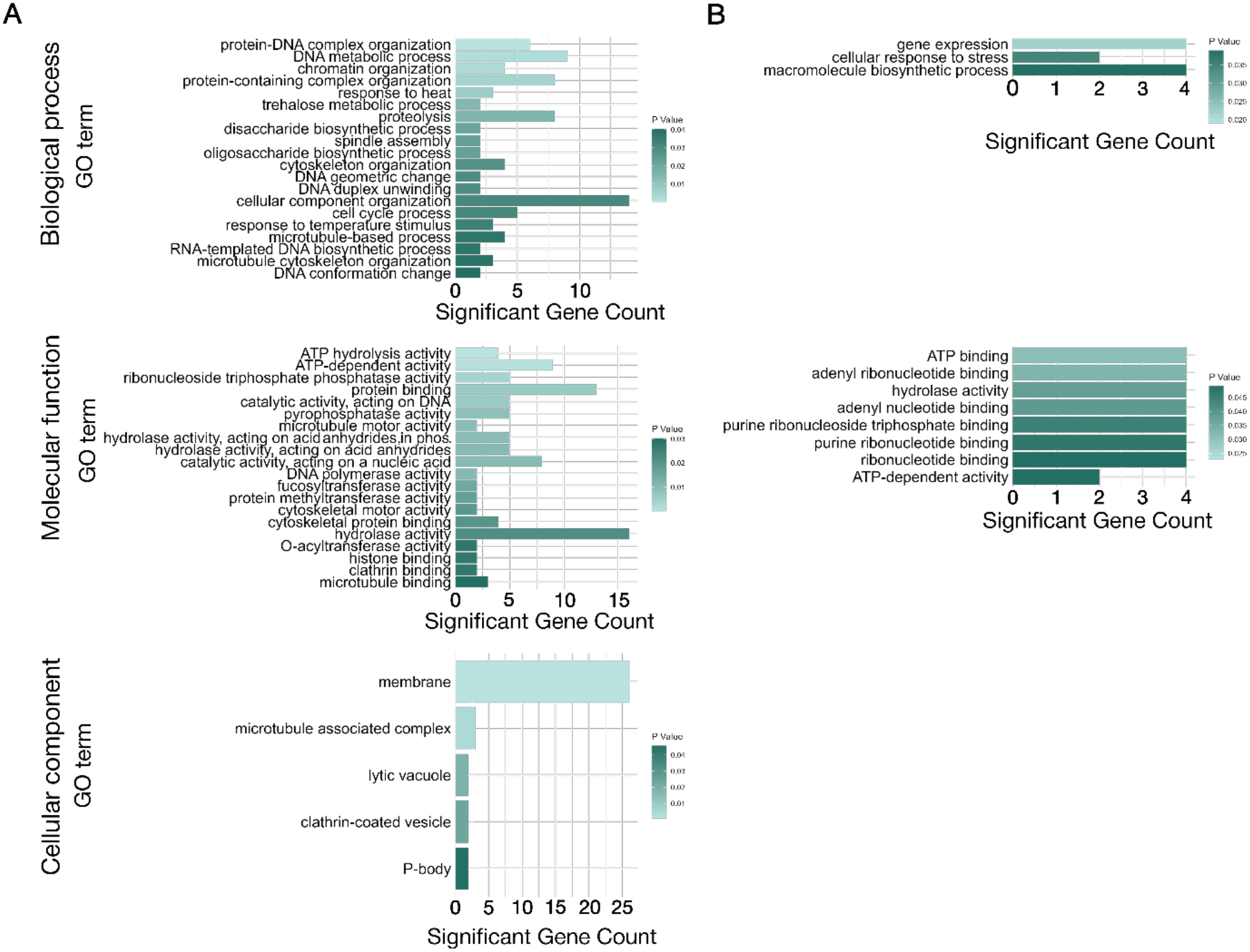
Gene set enrichment analysis of upregulated genes. **(A)** Gene ontology (GO) enrichment analysis of genes upregulated on the B chromosome. **(B)** GO enrichment analysis of genes upregulated on the A chromosomes. Bar plots show significantly enriched GO terms (*P* < 0.05) for molecular function, cellular component, and biological process categories. Bar length represents the number of upregulated genes associated with each GO term, while bar colour indicates the significance level (*P* value).

Among biological process terms, genes on chromosome B were significantly enriched for “chromatin organization” (GO:0006325), followed by “response to heat” (GO:0009408), “trehalose metabolic process” (GO:0005991), “proteolysis” (GO:0006508), “disaccharide biosynthetic process” (GO:0046351), “spindle assembly” (GO:0051225), “DNA duplex unwinding” (GO:0032508), and “RNA-templated DNA biosynthetic process” (GO:0006278). In the molecular function category, the most enriched GO term was “ATP hydrolysis activity” (GO:0016887), followed by “microtubule motor activity” (GO:0003777), “DNA polymerase activity” (GO:0034061), “fucosyltransferase activity” (GO: 0008417), “protein methyltransferase activity” (GO:0008276), “O-acyltransferase activity” (GO: 0008374), “histone binding” (GO:0042393), clathrin binding (GO: 0030276) and “microtubule binding” (GO:0008017). In the cellular component category, “membrane” (GO:0016020) was the most abundant GO term, followed by “microtubule associated complex” (GO:0005875), “lytic vacuole” (GO: 0000323), clathrin-coated vesicle (GO: 0030136) and “P-body” (GO:0000932) (Fig. 2A).

### ATAC-seq

We performed ATAC-seq on flow-sorted maize pollen nuclei (Supplementary Fig. S2) of the inbred line B73 plants (+B/0B), using three biological replicates per condition. Across all six libraries, read trimming retained nearly all reads (99.9 % on average; Supplementary Dataset S4), and the removal of mitochondrial and plastid DNA also preserved the majority of reads (96.15 % on average; Supplementary Dataset S5). Following downstream processing and filtering against a blacklist of highly repetitive regions and unplaced contigs, the final number of called peaks ranged from 71,184 to 80,052 per replicate; most libraries exhibited broadly comparable peak numbers, with a slightly higher count observed in +B replicates (Supplementary Dataset S5).

Overall, these data provided sufficient coverage to compare chromatin accessibility landscapes between pollen with and without B chromosomes (Cherchame *et al*., 2025). The consensus peak set consisted of 141,793 accessible chromatin regions (Supplementary Dataset S6). The genomic distribution of peaks across chromosomes was broadly similar among replicates for the standard A chromosomes (Supplementary Dataset S7), indicating that the core pollen accessibility landscape was largely shared between 0B and +B samples. As expected, a clear increase of peak number was observed for the B chromosome in +B replicates (Supplementary Dataset S7). In 0B samples, on average 334 peaks were assigned to the B chromosome due to mapping bias, whereas in +B samples chromosome B accounted for an average of 2,177 peaks (Supplementary Dataset S7). This pattern is consistent with the presence of B-chromosomal DNA only in the +B genotype and confirms that the B chromosomes maintain their own accessible chromatin landscape in pollen nuclei.

Comparison of peak sets between conditions revealed that the majority of accessible regions (72,375 peaks out of 141,793) were shared between 0B and +B pollen, which is consistent with the common cellular identity of the samples. When restricting the analysis to peaks uniquely associated with each condition, 31,803 peaks were specific to 0B and 37,615 peaks were specific to +B (Supplementary Figs. S3A; S3B). This difference indicates that the presence of B chromosomes is associated with an increased number of condition-specific accessible chromatin regions. Genomic annotation revealed that in both conditions (+B and 0B) the majority of peaks were located in distal intergenic regions (25,867-31,703), followed by promoter-proximal (24,509-27,770), intronic (1,589-2,326), and exonic (1,904-2,918) categories (Supplementary Table S2). Importantly, peaks were more abundant in +B samples than in 0B samples peaks across most annotation classes, including promoter-associated regions (4,472 for +B; 3,259 for 0B; Supplementary Table S3; Supplementary Table S4; Supplementary Fig. S4; Supplementary Fig. S5). This suggests that the presence of B chromosomes is associated not only with additional accessible intergenic regions, but also with accessibility changes near genes and putative regulatory elements. To further characterize the genomic context of accessible regions in the presence of the B chromosome, we examined the distribution of peaks across regulatory features in the +B condition. As expected, the majority of peaks (22,917) were located in distal intergenic regions; however, a substantial number of peaks on the B chromosome were also found in promoter-proximal regions (4,472), promoter regions (1-2 kb; 1,320), 5’ UTRs (247), 3’ UTRs (1,980), and gene bodies (3,042 exon and 2,633 intron peaks) (Fig. 3A).

**Figure 3.**
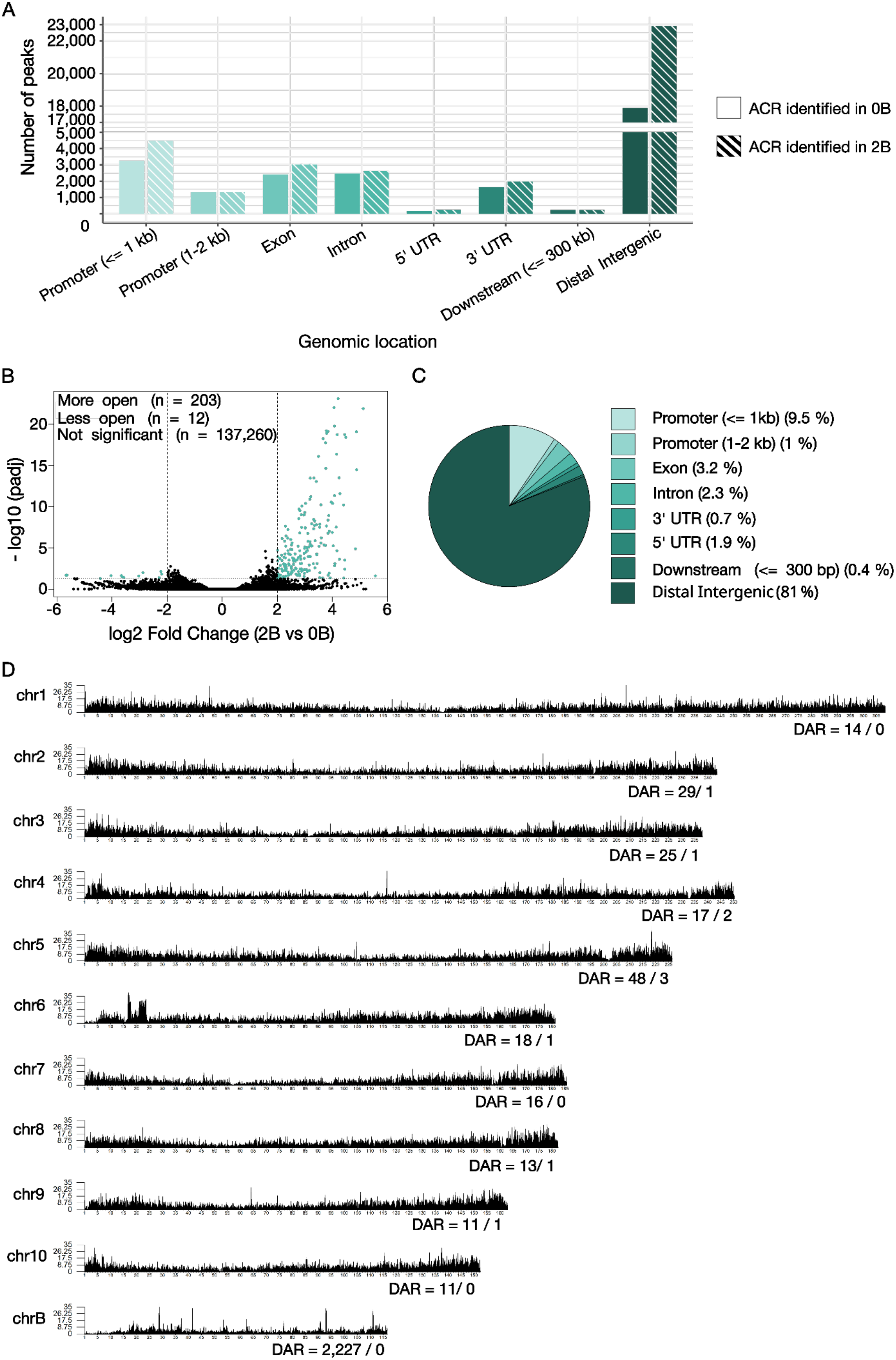
Distribution and genomic annotation of accessible chromatin regions and differentially accessible regions in maize pollen. **(A)** Distribution of accessible chromatin regions across the 0B/+B pollen nuclei and their genomic annotation. **(B)** Differential analysis of accessible chromatin regions on the A chromosomes. Volcano plot shows differential accessibility between +B and 0B pollen, with differentially accessible regions identified according to the applied significance and fold-change thresholds. The x-axis represents log2 fold change and the y-axis represents -log10 (adjusted *P* value). Dashed lines indicate the thresholds used to define differentially expressed genes. A total of 203 regions were more accessible and 12 regions were less accessible, while 137,260 regions were not significantly differentially accessible. **(C)** Genomic annotation of differentially more accessible chromatin regions (DARs) on the A chromosomes, showing their distribution among genomic features. **(D)** Distribution of DARs along chromosomes 1-10 and the B chromosome. Peak density is shown along each chromosome, and the numbers indicate the number of more/less differentially accessible regions identified on the respective chromosome.

### Differential analysis of accessible chromatin regions (ACRs)

Differential accessibility analysis identified 2,905 regions as significantly more accessible (203 on A chromosomes and 2,702 on the B chromosome) and 12 regions as significantly less accessible in the +B versus 0B comparison (Figs. 3B; 3C; 3D). Interpretation of chromosome assignment in this context must consider the fact that the B chromosome is absent from 0B plants. Therefore, peaks on the B chromosome can only contribute to the class of regions gaining accessibility in the presence of the B chromosome, whereas a category of “less accessible B-chromosome chromatin regions” lacks biological symmetry in this experimental design. With this limitation in mind, the data show that the vast majority of more accessible regions were mapped to the B chromosome (2,702 ACRs; Supplementary Dataset S8), while a smaller number of accessible peaks were located on A chromosomes (203 ACRs; Figs. 3B - 3D). Thus, the main accessibility gain associated with the presence of the B chromosome reflects the addition of accessible B-chromosomal regions. Genome-wide peak density analysis revealed localized regions of exceptionally high chromatin accessibility on chromosome 6 (Fig. 3D). Sequence annotation indicated that the region on chromosome 6 corresponds to ribosomal DNA (rDNA) clusters, consistent with known nucleolar organizer regions (NORs) in maize. Importantly, these regions did not exhibit strong differential accessibility between 0B and +B samples, indicating that they reflect constitutively accessible genomic features (Liu *et al*., 2025) rather than B chromosome-associated regulatory changes. More interestingly, in comparison with other A chromosomes, an increased number of differentially accessible chromatin regions were observed on chromosome 5 (Fig. 3D).

Genome-wide visualization of peak density revealed distinct differences between A and B chromosomes. When considering only differentially accessible regions (DARs), A chromosomes showed relatively sparse and dispersed changes, whereas the B chromosome displayed strong regional enrichment of peaks with increased accessibility in the +B condition. Although this strong enrichment on B chromosome was captured via DESeq2, statistically a high number of peaks were classified as significantly more open simply because they are missing in 0B samples. Nevertheless, these results indicate that B chromosomes behave as an accessible and dynamic chromatin feature.

ACRs, that were more accessible in samples with two B chromosomes, were associated with 738 nearby genes (168 genes on A chromosomes, after removal of duplicate peak-gene assignments), and a substantial fraction of these genes were B-linked (570 genes, after removal of duplicate peak-gene assignments; Supplementary Dataset S8). In contrast, peaks that were less accessible were associated exclusively with A-chromosome genes (12 genes). Because B chromosomes are absent from the 0B genotype, this asymmetry should not be interpreted simply as evidence that B-linked loci are preferentially protected from repression. Rather, it indicates that the presence of B chromosomes introduces an additional set of accessible loci, most of which are positioned near annotated B-linked genes, while reductions in accessibility are confined to loci present in both genotypes. However, the effect of the B chromosome on the accessibility of chromatin in the vicinity of A-chromosome-encoded genes should not be overlooked.

### Gene set enrichment analyis of genes near more open chromatin

We performed Gene Ontology (GO) enrichment analysis using the topGO package to investigate the functional roles of genes located near more open chromatin regions (168 genes on A chromosomes and 570 on the B chromosome). Where several significant terms were related, we reported the more specific term (i.e., the one lower in the GO hierarchy) over its more general parent term to maximize informativeness. The complete lists of tested GO terms, including non-significant ones, are provided in Supplementary Dataset S4. Among biological processes, genes on chromosome A were significantly enriched for “homeostatic process” (GO:0042592) followed by “rRNA processing” (GO:0006364). In the molecular function category, the most enriched GO term was “chromatin binding” (GO:0003682), followed by “transcription coregulator activity” (GO:0003712), and “enzyme binding” (GO:0019899). In contrast, no significantly enriched GO terms were detected within the cellular component category (Fig. 4B).

**Figure 4.**
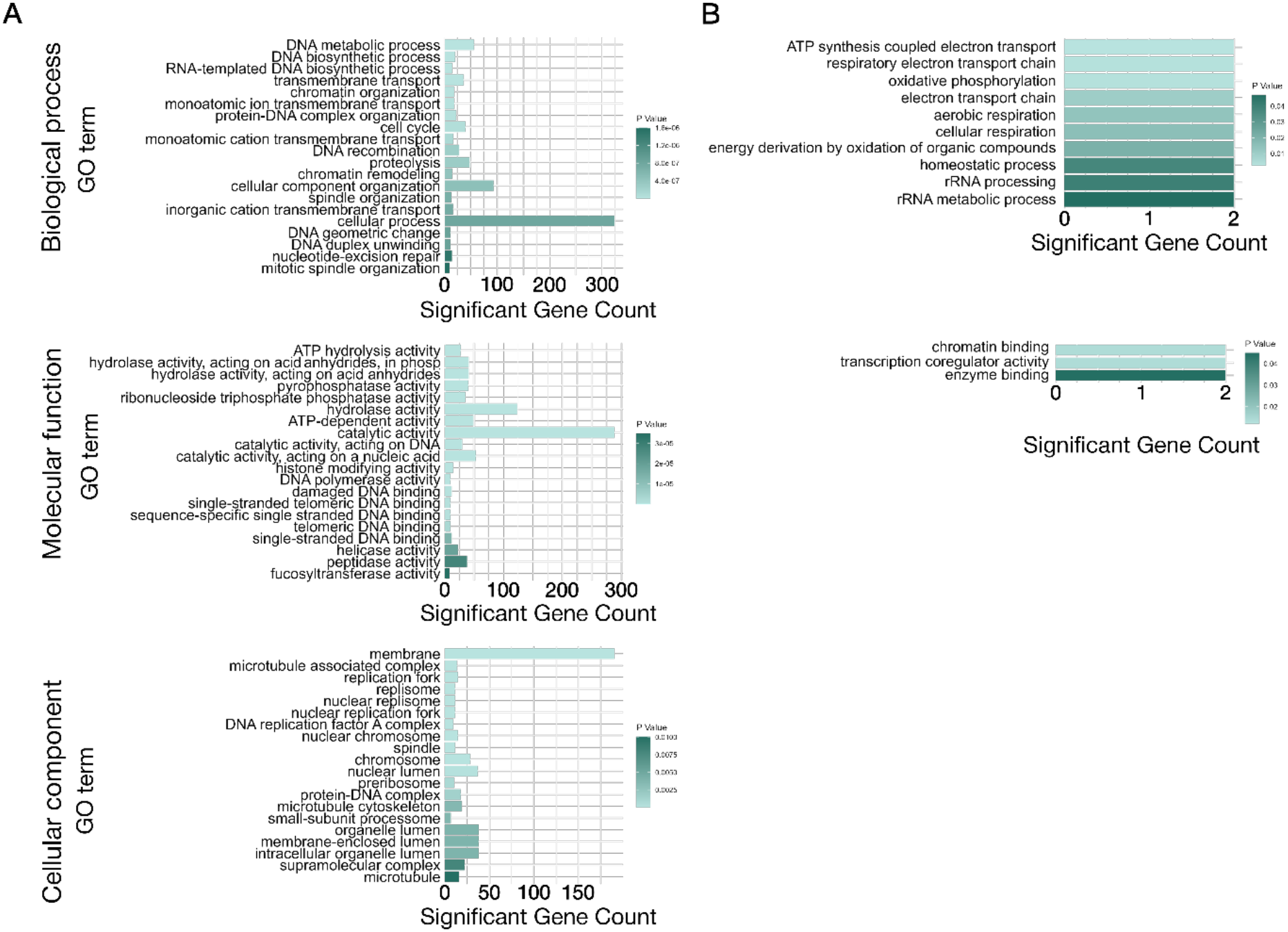
Gene set enrichment analysis of genes near more open chromatin regions. **(A)** Gene ontology (GO) enrichment analysis of genes near more open chromatin regions on the B chromosome. **(B)** GO enrichment analysis of genes near more open chromatin regions on the A chromosomes. Bar plots show significantly enriched GO terms (*P* < 0.05) for molecular function, cellular component, and biological process categories. Bar length represents the number of upregulated genes associated with each GO term, while bar colour indicates the significance level (*P* value).

For B-chromosome-located genes, the most prevalent biological process GO term was “RNA-templated DNA biosynthetic process” (GO:0006278), followed by “monoatomic cation transmembrane transport” (GO:0098655), “DNA recombination” (GO:0006310), “proteolysis” (GO: 0006508), “chromatin remodelling” (GO:0006338), “inorganic cation transmembrane transport” (GO:0098662), “DNA duplex unwinding” (GO:0032508), “nucleotide-excision repair” (GO:0006289) and “mitotic spindle organization” (GO:0007052). In the molecular function category, the most enriched GO term was “ATP hydrolysis activity” (GO:0016887), followed by “histone modifying activity” (GO:0140993), “DNA polymerase activity” (GO:0034061), “damaged DNA binding” (GO:0003684), “single-stranded telomeric DNA binding” (GO:0043047), “helicase activity” (GO:0004386), “peptidase activity” (GO:0008233) and “fucosyltransferase activity” (GO:0008417). Finally. Within the cellular component category, “membrane” (GO:0016020) was the most abundant GO term, followed by “microtubule associated complex” (GO:0005875), “DNA replication factor A complex”

(GO:0005662), “spindle” (GO:0005819), “small-subunit processome” (GO:0032040) and “microtubule” (GO:0005874) (Fig. 4A).

### Motif enrichment analysis

Transcription factors (TF) typically bind to short DNA sequences spanning 6-15 bp. We analysed motifs within the differentially accessible chromatin regions (DARs) to predict TFs that preferentially bind under the presence of B chromosomes. Motif enrichment analysis of the strongly enhanced ATAC-seq peaks (2,905 DARs; adjusted P < 0.05 and log2 fold change ≥ 2) identified several known motifs that were significantly enriched relative to the complete consensus peak set of 141,793 accessible regions (Table 1; Supplementary Dataset S10). Specifically, multiple independent TCP motifs showed a 1.3 to 2-fold enrichment (At1g69690/TCP: 1.77-fold; At2g45680/TCP: 1.95-fold; TCP20: 1.45-fold; At1g72010 TCP: 1.38-fold; At5g08330/TCP: 1.35-fold; TCP7: 1.38-fold), providing a consistent regulatory signature associate with the TCP-family, which is known to regulate reproductive development, cell cycle dynamics, proliferation, and hormonal pathways (Viola and Gonzalez, 2023). A second enriched group comprised MYB-related motifs (1.2-to 1.3-fold enrichment; involved in cell wall biosynthesis, stress responses, and hormone signalling; Wu *et al*., 2024). Additionally, significant enrichment of FHY3/FAR1-(1.27-fold), Trihelix-(1.1-to 1.3-fold), CAMTA-(1.2-fold), C3H-(1.49-fold), GATA/ZML-(1.63-to 1.96-fold) and E2F-like (1.2-to 3.9-fold) motifs.

**Table 1:**
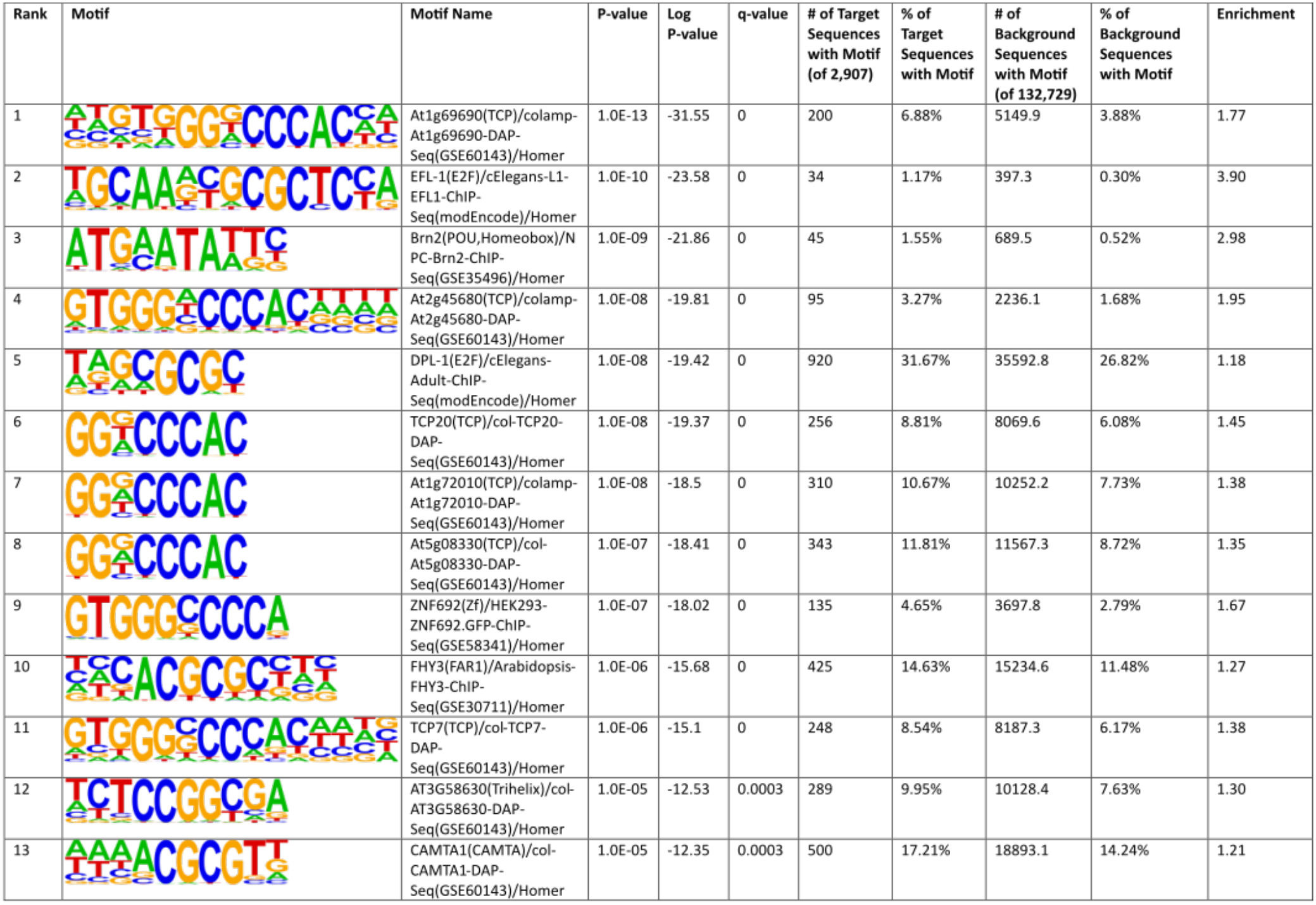
HOMER known motif enrichment analysis of ATAC-seq regions showing increased chromatin accessibility in the presence of B chromosomes. The 13 highest-ranked known motifs identified by HOMER are shown for regions with significantly increased accessibility. Motif occurrence in the target regions (n = 2,907) was compared with the background set of accessible chromatin regions (n = 132,729). Motifs are ranked according to enrichment *P*-value. Target and background frequencies indicate the proportion of sequences containing each motif, and enrichment was calculated as the ratio of target to background motif frequencies.

| Rank | Motif | Motif Name | P-value | Log P-value | q-value | # of Target Sequences with Motif (of 2,907) | % of Target Sequences with Motif | # of Background Sequences with Motif (of 132,729) | % of Background Sequences with Motif | Enrichment |
|---|---|---|---|---|---|---|---|---|---|---|
| 1 |  | At1g69690(TCP)/colamp-At1g69690-DAP-Seq(GSE60143)/Homer | 1.0E-13 | -31.55 | 0 | 200 | 6.88% | 5149.9 | 3.88% | 1.77 |
| 2 |  | EFL-1(E2F)/cElegans-L1-EFL1-ChIP-Seq(modEncode)/Homer | 1.0E-10 | -23.58 | 0 | 34 | 1.17% | 397.3 | 0.30% | 3.90 |
| 3 |  | Brn2(POU,Homeobox)/NPC-Brn2-ChIP-Seq(GSE35496)/Homer | 1.0E-09 | -21.86 | 0 | 45 | 1.55% | 689.5 | 0.52% | 2.98 |
| 4 |  | At2g45680(TCP)/colamp-At2g45680-DAP-Seq(GSE60143)/Homer | 1.0E-08 | -19.81 | 0 | 95 | 3.27% | 2236.1 | 1.68% | 1.95 |
| 5 |  | DPL-1(E2F)/cElegans-Adult-ChIP-Seq(modEncode)/Homer | 1.0E-08 | -19.42 | 0 | 920 | 31.67% | 35592.8 | 26.82% | 1.18 |
| 6 |  | TCP20(TCP)/col-TCP20-DAP-Seq(GSE60143)/Homer | 1.0E-08 | -19.37 | 0 | 256 | 8.81% | 8069.6 | 6.08% | 1.45 |
| 7 |  | At1g72010(TCP)/colamp-At1g72010-DAP-Seq(GSE60143)/Homer | 1.0E-08 | -18.5 | 0 | 310 | 10.67% | 10252.2 | 7.73% | 1.38 |
| 8 |  | At5g08330(TCP)/col-At5g08330-DAP-Seq(GSE60143)/Homer | 1.0E-07 | -18.41 | 0 | 343 | 11.81% | 11567.3 | 8.72% | 1.35 |
| 9 |  | ZNF692(Zf)/HEK293-ZNF692.GFP-ChIP-Seq(GSE58341)/Homer | 1.0E-07 | -18.02 | 0 | 135 | 4.65% | 3697.8 | 2.79% | 1.67 |
| 10 |  | FHY3(FAR1)/Arabidopsis-FHY3-ChIP-Seq(GSE30711)/Homer | 1.0E-06 | -15.68 | 0 | 425 | 14.63% | 15234.6 | 11.48% | 1.27 |
| 11 |  | TCP7(TCP)/col-TCP7-DAP-Seq(GSE60143)/Homer | 1.0E-06 | -15.1 | 0 | 248 | 8.54% | 8187.3 | 6.17% | 1.38 |
| 12 |  | AT3G58630(Trihelix)/col-AT3G58630-DAP-Seq(GSE60143)/Homer | 1.0E-05 | -12.53 | 0.0003 | 289 | 9.95% | 10128.4 | 7.63% | 1.30 |
| 13 |  | CAMTA1(CAMTA)/col-CAMTA1-DAP-Seq(GSE60143)/Homer | 1.0E-05 | -12.35 | 0.0003 | 500 | 17.21% | 18893.1 | 14.24% | 1.21 |

### Integration of ATAC-seq and RNA-seq

Numerous recent studies have demonstrated that the B chromosomes are not merely structurally present, but also transcriptionally active. In anthers containing mature pollen, hundreds of B-chromosome-encoded genes have been detected among expressed genes and among significantly upregulated genes (Hloušková *et al*., 2025). In our mature pollen RNA-seq dataset, the total number of expressed genes was lower, but B-linked genes still represented a substantial fraction of the significantly upregulated genes. To identify loci showing coordinated changes in chromatin accessibility and gene expression, we intersected the pollen ATAC-seq and pollen RNA-seq datasets (Fig. 5B). Importantly, this overlap identified 87 genes that exhibited both significantly increased chromatin accessibility and elevated transcript abundance, the vast majority of which (83) corresponded to B-linked loci (Fig. 5B). By contrast, no genes showed a reciprocal match between reduced accessibility and reduced expression.

**Figure 5.**
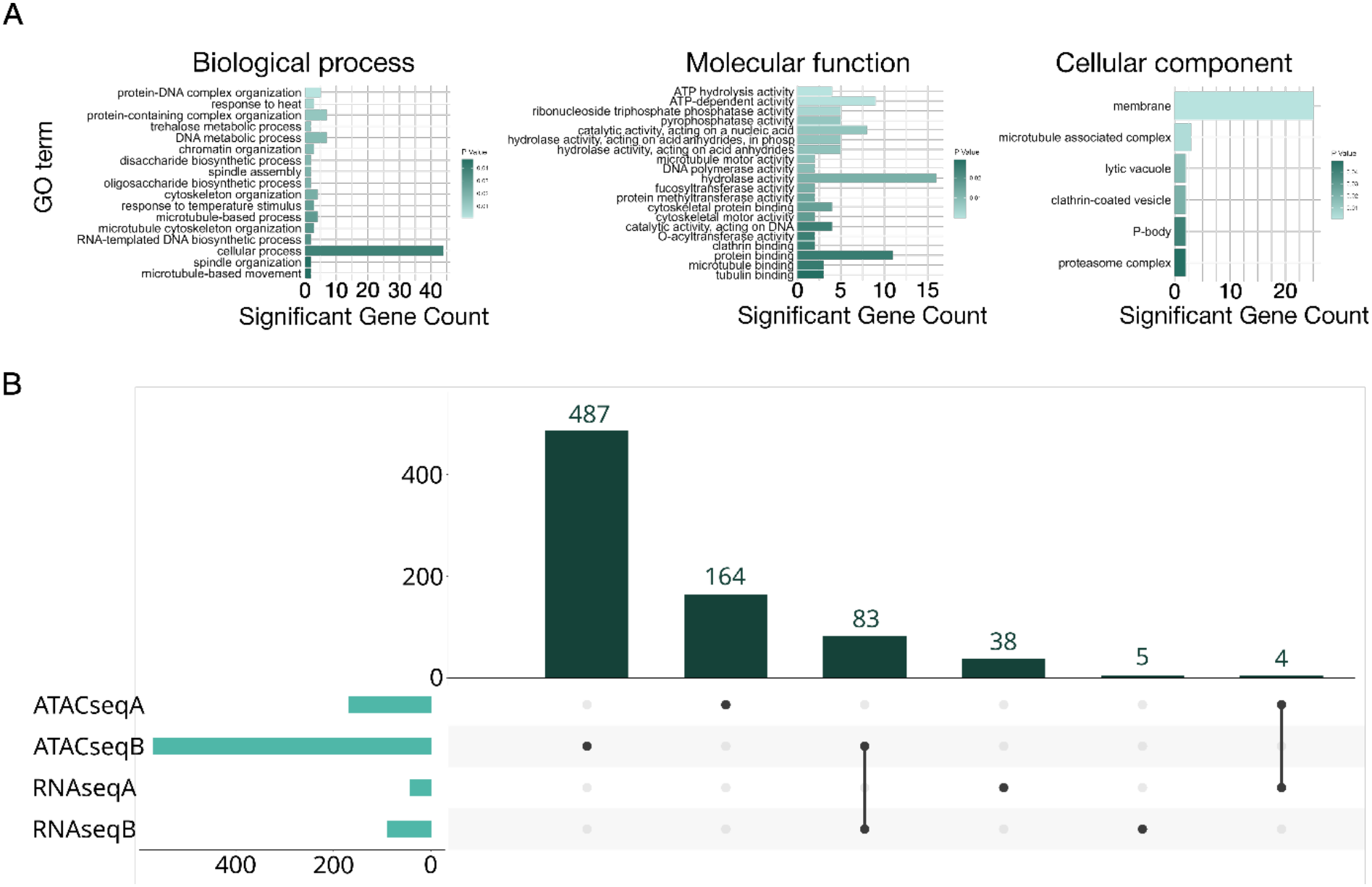
Gene set enrichment analysis of overlapping upregulated RNA-seq genes and genes near more open chromatin regions. **(A)** Gene ontology (GO) enrichment analysis of of overlapping upregulated RNA-seq genes and genes near more open chromatin regions. Bar plots show significantly enriched GO terms (*P* < 0.05) for molecular function, cellular component, and biological process categories. Bar length represents the number of upregulated genes associated with each GO term, while bar colour indicates the significance level (*P* value). **(B)** UpSet plot showing intersection of the genes located near the more open chromatin regions in the presence of the B chromosome (ATACseq) and genes upregulated in the presence of the B chromosome (RNAseq). The localization of genes on A chromosomes and B chromosome is indicated by corresponding suffix.

Together, these results support the conclusion that the presence of B chromosomes in maize pollen is associated with the establishment of accessible chromatin at B-linked loci and the concomitant transcriptional upregulation of a subset of B-chromosome genes.

At the same time, the data suggest that the influence of B chromosomes is not restricted entirely to the B genome itself. A portion of differentially accessible regions (DARs) and differentially expressed genes were assigned to A chromosomes, however, only four A-chromosome genes were specifically highlighted as showing concordant changes in both pollen ATAC-seq and pollen RNA-seq (Fig. 5B). These included loci associated with promoter-proximal as well as distal regulatory regions and encompassed genes with diverse predicted functions: RNA processing (*Zm00001eb048500*), helicase activity (*Zm00001eb164280*), lipid metabolism (*Zm00001eb224090*), and DNA replication/repair (*Zm00001eb330170*) (Supplementary Dataset S9).

To evaluate the functional profile of all 87 coregulated genes, we performed a GO enrichment analysis. Among biological processes, the most enriched was “response to heat” (GO:0009408), followed by “trehalose metabolic process” (GO:0005991), “chromatin organization” (GO:0006325), “disaccharide biosynthetic process” (GO:0046351), “spindle assembly” (GO:0051225) and “RNA-templated DNA biosynthetic process” (GO:0006278) and “microtubule-based movement” (GO:0007018). In the molecular function category, the most enriched GO term was “ATP hydrolysis activity” (GO:0016887), followed by “microtubule motor activity” (GO:0003777), “DNA polymerase activity” (GO:0034061), “fucosyltransferase activity” (GO:0008417), “protein methyltransferase activity” (GO:0008276), “O-acyltransferase activity” (GO:0008374), “clathrin binding” (GO:0030276) and “microtubule binding” (GO:0008017). Within the cellular component category, “membrane” (GO:0016020) was the most abundant GO term, followed by “microtubule associated complex” (GO:0005875), “lytic vacuole” (GO:0000323), “clathrin-coated vesicle” (GO:0030136), “P-body” (GO:0000932) and “proteasome complex” (GO:0000502) (Fig. 5B). Where several significant terms were related, we reported the more specific term (i.e., the one lower in the GO hierarchy) over its more general parent term to maximize informativeness. The complete lists of tested GO terms, including non-significant ones, are provided in Supplementary Dataset S4.

These observations indicate that the presence of B chromosomes may also have secondary effects on the regulatory state of selected A-chromosome loci. However, this effect appears limited compared with the much stronger signal generated directly by B-linked accessible chromatin and transcription. Overall, the results indicate that maize B chromosomes in pollen are associated with a distinct accessible chromatin profile and with transcriptional activity, particularly at B-linked loci.

The ATAC-seq and RNA-seq datasets capture different levels of the regulatory response (Fig. 5B; Supplementary Dataset S9). Specifically, differentially accessible regions show a strong enrichment for TCP-family signatures (Table 1; Supplementary Dataset S10), whereas the most strongly induced transcripts are dominated by stress-and proteostasis-related functions, RNA metabolism, and several chromatin or transcription-associated factors (Supplementary Dataset S9). Intriguingly, the only strongly induced, sequence-specific TF within the filtered A-chromosome DEG set is an AP2/ERF-family factor (Zm00001eb307550/ERF1B), whose motif family is not significantly enriched among the opening chromatin regions (Supplementary Dataset S9; Supplementary Dataset S10).

## Discussion

We provide the first evidence that maize B chromosomes possess an accessible regulatory landscape that is coupled to active transcription in mature pollen. While previous work has extensively described the mechanism of B chromosome transmission through chromosome drive, little was known about their chromatin organization and transcriptional activity in the male gametophyte. By combining flow-sorted pollen nuclei with ATAC-seq and RNA-seq, we demonstrate that B chromosomes harbour numerous accessible chromatin regions and actively transcribed genes, indicating that they are functionally active components of the pollen genome rather than genetically inert elements. Because the B chromosome is physically absent from 0B plants, the strongest conclusion is not that chromatin accessibility is globally redistributed between equivalent genomic compartments, but rather that the addition of B chromosomes introduces a new set of accessible and transcriptionally active regions into the pollen nucleus. Together, these findings indicate that such activity is predominantly restricted to the B chromosome itself, as its influence on the standard A genome appears comparatively modest.

While mature maize pollen grains show an overall lower gene expression level and a reduced number of expressed genes (TPM > 1; Shi *et al*., 2021; Han *et al*., 2022; Nelms and Walbot, 2022; Zhang *et al*., 2022; Hloušková *et al*., 2025) relative to anther and other tissues, they simultaneously maintain a subset of distinct, highly enriched pollen-specific genes (Honys and Twell, 2003; Pina *et al*., 2005; Ma *et al*., 2006; Ling *et al*., 2015; Shi *et al*., 2021; Hong *et al*., 2022; Le Lievre *et al*., 2023; Supplementary Dataset S2). The less complex pollen transcriptome observed here is consistent with previous studies and represents a conserved feature among diverse species (Honys and Twell, 2003; Gómez *et al*., 2015; Han *et al*., 2022). Expression profiling of tissues from B-positive plants revealed that 103 of the 689 predicted B-chromosome-encoded genes were transcribed in mature pollen (TPM > 1; Supplementary Dataset S1), with 98 of these genes overlapping with those detected in the whole anther transcriptome. In contrast to our results, Hloušková *et al*. (2025) previously reported that 352 B chromosome-located genes are actively transcribed across a wider selection of tissues including 244 expressed B-genes in anther tissue containing mature pollen. Combined with our detection of B-encoded genes in mature pollen, these data strongly suggest that B chromosomes are preferentially expressed in reproductive organs, implying a potential functional role for their genes in reproductive development, maturation, and transmission of the B chromosome to the next generation.

The detection of substantial amounts of B-chromosome mRNA in mature pollen further suggests that these transcripts are stored for subsequent use during pollen germination and tube growth as previously described by Rutley and Twell (2015). The germination and guided growth of pollen tubes are driven by a combination of stored transcripts and newly synthesized mRNAs that facilitate signalling and maternal-tissue communication within the gynoecium (Rutley and Twell, 2015). Consistent with the fundamental importance of pollen-pistil communication, the pollen transcriptome was highly enriched for signalling components known to facilitate female gametophyte-driven guidance of the pollen tube to the ovule (Palanivelu and Tsukamoto, 2012). Conversely, processes related to energy metabolism, transport, transcription, and translation remain under-represented (Rutley and Twell, 2015). Compared to sporophytic tissues, the maize pollen transcriptome exhibits reduced total gene diversity alongside a highly skewed expression profile characterized by exceptionally high maximum transcript levels (Davidson *et al*., 2011).

Critical to the continuation of the B chromosome’s survival in the genetic lineage of maize is its drive mechanism. While nondisjunction at the second pollen mitosis is controlled by cis and trans factors on the B chromosome (Birchler and Yang 2021; Blavet *et al*., 2021) and occurs when sister chromatids fail to separate at the centromeric region, the second component of the drive is arguably more relevant to the biology of mature pollen. Current evidence suggests this mechanism involves structural domains adjacent to the B centromere-proximal region alongside egg cell-driven recognition of specific sperm nuclei, although the precise molecular factors have yet to be identified (González-Sánchez *et al*., 2003; Brennan *et al*., 2023).

The B chromosome is distinguished by repeat-dense pericentromeric regions, notably ZmBs repeats and knob-like heterochromatin (Birchler and Yang, 2021; Blavet *et al*., 2021). Within the sperm nucleus, these B-specific sequences adopt a localized spatial organization at one pole or subregion, imparting a distinct nuclear architecture that may influence gametic recognition during double fertilization (Rusche *et al*., 1997; Blavet *et al*., 2021). When the egg and central cell are both accessible to the two sperm, the egg appears to differentially interact with the B-carrying sperm nucleus, likely via specific receptor-ligand, adhesion, or signalling components on the egg plasma membrane or its surroundings. This mechanism may involve egg-cell-expressed proteins or RNAs that recognize features associated with the B-centromere region, such as centromeric proteins, unique chromatin marks, or nuclear envelope contacts, leading to more efficient fusion or nuclear migration. Furthermore, since the B centromere region is enriched in specific repeats related to knob heterochromatin, these regions replicate late and can alter chromatin compaction or nuclear architecture, potentially changing how the sperm nucleus interacts with the egg-cell nuclear envelope or cytoskeletal guidance. A feasible, though not yet directly demonstrated, model is that egg-cell factors recognize nuclear envelope or centromere/kinetochore proteins associated with the B region, allowing the B-carrying sperm nucleus to be preferentially anchored, imported, or fused within the egg, while the other sperm nucleus associates with the central cell (González-Sánchez *et al*., 2003; Blavet *et al*., 2021).

Alternatively, preferential fertilization might initiate earlier, within the pollen tube, if the B-carrying sperm exhibits altered interactions with the vegetative cell or tube cytoskeleton that position it closer to the egg at discharge, or affect the sequence in which the two sperm are released and fuse. B-linked factors could modulate the expression or localization of proteins in the generative or sperm cells that influence motility-like behaviours inside the tube, such as movement along actin filaments, thereby giving the B-rich sperm a kinetic advantage toward the egg rather than the central cell (González-Sánchez *et al*., 2003; Blavet *et al*., 2021). Despite cytological and genetic evidence pointing toward a model governed by nuclear architecture and egg-cell recognition, the precise molecular machinery remains unknown. Specifically, the genes, proteins, and direct interactions at the egg membrane, nuclear envelope, or pollen tube that physically execute this preferential fertilization warrant further investigation.

Our results of differentially accessible chromatin extend observations of Hloušková *et al*., 2025 by demonstrating that B chromosome transcription is maintained in mature pollen -a highly specialized cell type characterized by a relatively simple transcriptome, as described in *A. thaliana* (Honys and Twell, 2003; Pina *et al*., 2005), rice (Wei *et al*., 2010; Hong *et al*., 2022), maize (Ma *et al*., 2006; Han *et al*., 2022) and cotton (Zhang *et al*., 2022). Typically, mRNA and rRNA gene transcripts accumulate and transcription is subsequently repressed during pollen maturation, with these transcripts stored for use in protein synthesis during pollen germination (Mascarenhas, 1989; Schrauwen *et al*., 1990; Honys *et al*., 2000; Honys and Twell 2003).

Although little is known about the higher-order chromatin organization of maize B chromosomes, recent work in other organisms suggests that supernumerary chromosomes can influence nuclear architecture. A recent chromosome-level assembly of the cichlid fish *Astatotilapia latifasciata* (Jehangir *et al*., 2026) revealed that a 34 Mb region of B chromosome acts as a major driver of nuclear architecture by reshaping chromatin organization. The presence of the B chromosome in this species is associated with fewer topologically associating domains (TADs), reduced chromatin loop formation, and altered genomic compartmentalization, while B-chromosome protein-coding genes are predominantly repressed relative to their A-chromosome paralogs. Furthermore, the B chromosome of the cichlid fish influences long-range chromatin interactions and is linked to genomic rearrangements within the host’s standard A chromosomes (Jehangir *et al*., 2026). Similarly, in wheat and rye, approximately 5-6% of A-derived transcripts were differentially expressed when two Bs were present (Boudichevskaia *et al*., 2022). In maize, the presence of B chromosomes has been shown to significantly alter the expression of over 3,000 genes located on A chromosomes, as well as small RNAs (miRNAs; Shi *et al*., 2022; Hloušková *et al*., 2025). Whether a gene is upregulated or downregulated often depends on the host species and tissue type (Boudichevskaia *et al*., 2022).

Interestingly, several loci exhibited accessible chromatin in both genotypes but increased transcript abundance only in the presence of B chromosomes. This observation suggests that chromatin accessibility alone is insufficient to explain transcriptional activation and that additional epigenetic or transcriptional regulatory mechanisms may contribute. Because B chromosomes depend on preferential transmission for their long-term persistence, selection may favour the retention of genes that enhance chromosome drive or ensure stable inheritance, while genes unrelated to these processes may gradually degenerate. It follows that candidate genes for the control of preferential fertilisation should be recruited from the list of B-chromosome-encoded genes expressed in mature and/or active pollen. Alternatively, some transcripts may simply result from the presence of accessible chromatin without conferring an immediate functional advantage. Further functional characterization of the expressed B-chromosome genes, particularly those associated with accessible promoter regions (232 genes out of 540 B genes; Supplementary Dataset S7), will be necessary to distinguish between these possibilities.

Beyond its biological findings, the present study demonstrates that flow cytometric purification of mature pollen nuclei can be successfully combined with ATAC-seq and RNA-seq to generate high-quality chromatin accessibility profiles. Mature pollen presents unique technical challenges because of its highly condensed chromatin and resistant pollen wall, making genome-wide epigenomic analyses difficult. Our workflow therefore provides a useful framework for future studies investigating chromatin organization in the male gametophyte.

Collectively, our findings demonstrate that maize B chromosomes establish a distinct chromatin accessibility landscape and contain transcriptionally active loci in mature pollen. Furthermore, the enrichment of accessible chromatin near B-linked genes suggests that regulatory activity is organized rather than randomly distributed across the B chromosome. Ultimately, this structural and transcriptional characterization provides new insight into the molecular organization of selfish chromosomes and establishes a foundation for future studies aimed at identifying the molecular mechanisms underlying B-chromosome maintenance, chromosome drive, and interactions with the host genome.

## Materials and Methods

### Plant Material and Growing Conditions

Mature pollen grains from maize (*Zea mays* L.) lines (B73 without the B chromosome - 0B; B73 possessing two B chromosomes - +B) were harvested from greenhouse-grown plants into tubes. The original seed stock of reference inbred line B73 was provided by the Plant Introduction Station (North Central Regional Plant Introduction Station, Iowa, USA) and the original seed stock of +B line by prof. Birchler (University of Missouri, Missouri, USA). Seeds were germinated at 25°C for 5 days, selected plants were grown in the greenhouse under controlled conditions of 14h of light at 24°C and 10h of dark at 20°C. A direct PCR approach was used to assess the presence of the B chromosome in individual seedlings. The +B seedlings were further scored using droplet digital PCR (ddPCR) protocol to estimate the number of B chromosomes (Svačina *et al*., 2023). Mature pollen (in three biological replicates for both - 0B and +B line) was immediately used for nuclei isolation and subsequent fluorescence-activated nuclei sorting by flow cytometry.

### Nuclei isolation and Flow cytometry

Anthers with mature pollen grains were vortexed in LB01 buffer (Doležel *et al*., 1989). Isolation was performed by bursting method according to Kron and Husband (2015). Suspension of pollen and anthers was filtered through a 100 μm filter (Celltrics, Sysmex) to remove anthers and through a 20 μm filter to capture pollen grains. The surface of the 20 μm filter with pollen was then gently rubbed with glass rod for a few seconds to release nuclei from pollen grains by washing down through the filter with LB01 buffer. Nuclei were stained with 4’,6-diamidino- 2-phenylindole (DAPI; 2 μg/ml) and loaded onto FACSAria II SORP flow cytometer and sorter (Becton Dickinson, San Jose, USA) equipped with 390 nm laser for DAPI excitation. Sorting gates were initially drawn on a dot plot of forward scattering (y-axis) and relative DAPI fluorescence (x-axis) and subsequently on a dot plot of relative DAPI-W fluorescence (y-axis) versus relative DAPI-A fluorescence (x-axis) to purify nuclei suspension. The individual fraction of 50,000 of all pollen (vegetative and sperm) nuclei was flow-sorted for each sample into LB01 with protease inhibitor (cOmplete, Roche) for subsequent ATAC-seq analysis (Grandi *et al*., 2022).

### ATAC-seq library construction and sequencing

Isolated nuclei were spun down at 500 x g for 5 minutes at 4 °C. During this time, tagmentation master mix was prepared with the ATAC-Seq Kit according to the manufacturer’s manual (Active Motif catalog No. 53150) to perform the tagmentation reaction that was held at 37 °C for 30 min in a thermal shaker (Eppendorf® ThermoMixer® C) at 750 rpm. Tagmented DNA was purified on purification column and PCR reaction for library amplification was performed according to the manufacturer protocol. The libraries were size selected using SPRI beads. DNA was quantified by Qubit HS DNA kit (Thermo Fisher, Q32854) and analyzed on Agilent 2100 BioAnalyzer for quality control. ATAC-seq libraries were sequenced on an Illumina NovaSeq 6000 platform (Illumina, San Diego, USA) which produced an average 200 million paired-end reads per sample (2×150 bp paired-end reads).

### RNA Extraction and Sequencing

Total RNA was isolated and sequenced for transcriptomic analysis according to Hloušková *et al*. (2025). RNA was isolated from pollen, in three biological replicates for reference B73 inbred line without B chromosomes and three biological replicas for B73 inbred line containing two B chromosomes, following the instruction provided in Monarch® Total RNA Miniprep Kit (New England Biolabs, Ipswich, USA). The quality of total RNA was evaluated using Agilent 2100 Bioanalyzer and the indexed sequencing libraries construction was performed according to the manufacturer’s protocol with NEBnext Ultra II RNA Library Prep Kit for Illumina (New England Biolabs). Libraries were sequenced on an Illumina NovaSeq 6000 platform, using 2×150-bp paired-end reads configuration, yielding ∼35 million paired-end reads per sample on average.

### ATAC-seq analysis

The analysis of ATAC-seq data was performed in Miniconda (v24.4.0; https://github.com/conda/conda), a cross-platform, open-source package and environment management system, to ensure compatibility among the tools and data formats. First, FastQC (v0.12.1; https://www.bioinformatics.babraham.ac.uk/projects/fastqc/) and MultiQC (v. 1.12; Ewels *et al*., 2016) were used for the quality control of the raw sequencing data to ensure that only high-quality reads are used in downstream analysis. The raw sequencing data were further trimmed using trim galore (v0.6.2; https://github.com/FelixKrueger/TrimGalore), and trimmed data were mapped to the *Zea may*s reference genome RefGen_v5 (Hufford *et al*., 2021) supplemented with the reference of the maize B chromosome (Maize_Bchr_v2.0, Hloušková *et al*., 2025), using a sensitive Bowtie2 v2.4.2 aligner (Langmead and Salzberg, 2012). All reads mapping to the mitochondrial and plastid DNA were removed from the .bam files for subsequent analysis. Reads were sorted and indexed. Properly-paired end reads were selected for next analysis using -f 3 parameter in samtools view (SAMtools v1.11; Li *et al*., 2009). PCR duplicates were removed by MarkDuplicates function in Picard (v2.9; https://broadinstitute.github.io/picard/). Due to library size normalization, equivalently 27 millions of reads were then randomly selected by samtools view from each library (to achieve size of the smallest available library - 27,298,261 reads; Supplementary Dataset S4) and equivalently more reads (+ 6.2 %) were randomly selected for +B sample to compensate for the B chromosome size. Before the subsequent analysis, .bam files were converted to .bedpe files by bedtools bamtobed v2.26.0 (Quinlan, 2014). Significant broad peaks were then called using MACS2 v2.2.6 (Zhang *et al*., 2008). To retrieve the black list of repetitive regions, RepeatMasker (Smit, AFA, Hubley, R & Green, P. RepeatMasker Open-4.0. 2013-2015 <http://www.repeatmasker.org>) was used. For masking we used Maize TE Consortium (MTEC) curated library of repeats (https://github.com/oushujun/MTEC) and custom-made library of repeats of B chromosome sequence generated using DANTE and DANTE_LTR (Novák *et al*., 2024). Peaks overlapping the blacklist of highly repetitive regions were filtered out using bedtools intersect v.2.26.0 (Quinlan, 2014). Finally, annotation of accessible chromatin regions and peak-gene assignment were done using ChIPseeker v1.48.0 (Yu *et al*., 2015) and differential accessibility analysis was done using DESeq2 (v.1.12.3) package in R (RStudio (v3.4.0), Love *et al*., 2014) with *P*-value < 0.05 and Log_2_FoldChange > 2 or < -2 to identify significant differentially accessible chromatin regions. For manual control, data were visualized by Integrative Genomics Viewer (IGV; v2.17.4.; Robinson *et al*., 2011)

### Motif enrichment analysis

Motif enrichment analysis of differentially accessible ATAC-seq regions was performed using HOMER v.5.1 (Heinz *et al*., 2010). Regions with significantly increased accessibility (adjusted *P* < 0.05 and log2 fold change ≥ 2) were analyzed with findMotifsGenome.pl using the reference genome of *Zea mays* reference RefGen_v5 (Hufford *et al*., 2021) supplemented with the reference of the maize B chromosome (Maize_Bchr_v2.0, Hloušková *et al*., 2025). Motif searches were performed within 200-bp regions centered on each peak, testing motif lengths of 6, 8, 10, and 12 bp. The complete set of 141,794 ATAC-seq peaks was used as the background to control for sequence characteristics of accessible chromatin. Overlap of significant peaks with the complete ATAC peak set was verified using bedtools intersect.

### RNA-seq analysis

All RNA-seq reads were quality checked with FastQC (v.0.12.1; Andrews, 2010; https://www.bioinformatics.babraham.ac.uk/projects/fastqc/). The fastp software (Chen *et al*., 2018) was used to filter out sequencing adaptors and trim low-quality sequences (parameters: -q 30, -w 4, -l 50). Trimmed reads were mapped using STAR (Dobin *et al*., 2013) to the reference genome of *Zea mays* reference RefGen_v5 (Hufford *et al*., 2021) supplemented with the reference of the maize B chromosome (Maize_Bchr_v2.0, Hloušková *et al*., 2025). Mapped reads were counted using RSEM (Li and Dewey, 2011). Transcripts per Million (TPM) values were calculated using Tximport (v.1.34.0) package in R (Soneson *et al*., 2015). Differential-expression analysis was performed using DESeq2 (v.1.12.3) package in R (Love *et al*., 2014) with *P*-value < 0.05 and Log2FoldChange > 2 or < -2 to identify significant differentially expressed genes.

### Gene Set Enrichment Analysis

Gene set enrichment analysis (GSEA) was performed according to the Hloušková *et al*. (2025). Gene ontology (GO) terms provided for *Zea mays* reference RefGen_v5 (Hufford *et al*., 2021) were downloaded from MaizeGDB (https://www.maizegdb.org/). For the B chromosome-encoded genes, GO terms were obtained using Blast2GO pipeline (Hloušková *et al*., 2025). TopGO (v.2.58.0) package in R (Alexa and Rahnenfuhrer, 2024) was used to run a gene set enrichment analysis. GO terms were considered significantly enriched if the *P*-value from Fisher’s exact test was below 0.05 and the term was represented by more than one gene. GSEA was performed on subset of differentially upregulated genes located on A chromosomes and B chromosome-located genes and genes located near significantly more open chromatin regions on A chromosomes and the B chromosome and finally on all genes that overlapped between RNA-seq and genes near accessible chromatin region of ATAC-seq.

## Supplementary data

Supplementary Table S1. Summary statistics of Illumina NovaSeq transcriptome sequencing

Supplementary Table S2. Distribution of accessible chromatin regions among replicates

Supplementary Table S3. Distribution of accessible chromatin regions uniquely in 0B/+B conditions

Supplementary Table S4. Distribution of accessible chromatin regions including common regions for 0B/+B condition

Supplementary Figure S1. Chromosomal distribution of differentially expressed genes in maize anthers with pollen

Supplementary Figure S2. Flow cytometric analysis of pollen nuclei from maize inbred line B73 with and without B chromosomes

Supplementary Figure S3. Distribution of accessible chromatin regions (ACR) after DESeq2 among replicates.

Supplementary Figure S4. Genomic annotation of accessible chromatin regions (ACR) uniquely in 0B (A) and +B (B) condition on chromosomes.

Supplementary Figure S5. Genomic annotation of accessible chromatin regions (ACR) including common regions for 0B (A) and +B (B) condition on chromosomes.

Supplementary Dataset S1. TPM values of genes in 12 maize plant tissues in +B/0B conditions (genes expressed predominantly in pollen)

Supplementary Dataset S2. TPM values assigned for chromosome-encoded genes (the green shading indicates differences in expression)

Supplementary Dataset S3. Results of differential gene expression analysis in the pollen tissue in 0B/+B condition

Supplementary Dataset S4. Summary statistics of gene ontology enrichment analysis Supplementary Dataset S5. Summary statistics of reads and peaks in ATAC-seq workflow Supplementary Dataset S6. ATAC-seq consensus peak set

Supplementary Dataset S7. Distribution of accessible chromatin regions on chromosomes

Supplementary Dataset S8. Results of differential chromatin accessibility analysis in the pollen nuclei in 0B/+B condition

Supplementary Dataset S9. Gene description of differential gene expression analysis (RNA-seq), genes near differentially accessible chromatin analysis (ATAC-seq) and genes overlapping both methods

Supplementary Dataset S10. Homer known Motif enrichment results

All supplementary files are available at: https://zenodo.org/records/24077189?preview=1&token=eyJhbGciOiJIUzUxMiIsImlhdCI6MT c5NDIxNTU1MywiZXhwIjoxNzk4Njc1MTk5fQ.eyJpZCI6IjZmM2RhMmZiLWJlYWItNGFlYy04Nm FmLTM3YjQwOTBiMzlmNSIsImRhdGEiOnt9LCJyYW5kb20iOiI3OTkwNzNhYmY2YjU5MDJjNmE 1YzljMzkzOTc4OWQ2OSJ9.ZapwDv8MTpmCU_9vYlCWhnYRxjHhpesXM2D_VP-eSvVG6-4JCqQclY0peC51dve2em-v8FA14Z5EyT5jX3wHPw

## Acknowledgements

We thank Jitka Weisserová, Helena Tvardíková and Magdaléna Stejskalová for technical assistance. Computational resources were provided by the e-INFRA CZ project (ID:90254), supported by the Ministry of Education, Youth and Sports of the Czech Republic.

## Funding

This work was supported by the Czech Science Foundation (project no. 23-04887S) and by the Operation Program Research, Development, and Education project: Plants as a tool for sustainable global development (No.CZ.02.1.01/0.0/0.0/16_019/0000827).

## Author contributions

PU, JB – investigation; PU, LH, KH, JB – formal analysis; PU – writing – original draft; PU, LH, KH, JD, JB – review and editing; JB – conceptualization, project administration and funding acquisition.

## Competing interests

The authors declare they have no competing interests.

The authors declare that the research was conducted in the absence of any commercial or financial relationships that could be construed as a potential conflict of interest.

## Data availability

Raw sequence reads are available at the European Nucleotide Archive (https://www.ebi.ac.uk/ena/browser/home) under accession number PRJEB129204 (ATAC-seq and RNA-seq).

## Notes

### Competing Interest Statement

The authors have declared no competing interest.

